# *Song Torrent*: A modular, open-source 96-chamber audio and video recording apparatus with optogenetic activation and inactivation capabilities for *Drosophila*

**DOI:** 10.1101/2024.01.09.574712

**Authors:** Steve Sawtelle, Lakshmi Narayan, Yun Ding, Elizabeth Kim, Emily L. Behrman, Joshua L. Lillvis, Takashi Kawase, David L. Stern

## Abstract

**Background:**

- Many *Drosophila* species use acoustic communication during courtship and studies of these communication systems have provided insight into neurobiology, behavioral ecology, ethology, and evolution.
- Recording *Drosophila* courtship sounds and associated behavior is challenging, especially at high throughput, and previously designed devices are relatively expensive and complex to assemble.

**Results:**

- We present construction plans for a modular system utilizing mostly off-the-shelf, relatively inexpensive components that provides simultaneous high-resolution audio and video recording of 96 isolated or paired *Drosophila* individuals.
- We provide open-source control software to record audio and video.
- We designed high intensity LED arrays that can be used to perform optogenetic activation and inactivation of labelled neurons.
- The basic design can be modified to facilitate novel study designs or to record insects larger than *Drosophila*.
- Fewer than 96 microphones can be used in the system if the full array is not required or to reduce costs.

**Implications:**

- Our hardware design and software provide an improved platform for reliable and comparatively inexpensive high-throughput recording of *Drosophila* courtship acoustic and visual behavior and perhaps for recording acoustic signals of other small animals.

## Background

*Drosophila* flies perform elaborate social behaviors as part of courtship and mating, which in many species can include acoustic signals (Ewing and Bennet-Clark 1968). For example, depending on the species, males and/or females can produce complex and time-varying acoustic signals during courtship (LaRue *et al*. 2015; Arthur *et al*. 2021). In addition, males and females produce sounds during other social encounters, such as aggression and copulation (Versteven *et al*. 2017; Kerwin *et al*. 2020). Individual flies are highly motivated to perform these complex behaviors, even in artificial laboratory conditions, which has made *Drosophila* courtship behavior a powerful model for studies of the genetic and neurobiological basis of social behavior and evolution (Auer and Benton 2016).

The acoustic signals produced by *Drosophila* during social interactions are relatively weak and are best detected with particle-velocity sensitive microphones located close to flies (Bennet-Clark 1971, 1984) because the power of the particle velocity declines as the cube of the distance. This constraint has complicated the development of high-throughput recording devices of *Drosophila* courtship song. *Drosophila* courtship song has been studied by many laboratories over the past 60 years and has historically been recorded using bespoke amplifier and microphone apparatuses and analysis techniques. In 2013, we introduced a multi-channel device that allowed simultaneous recording of courtship song from 36 pairs of flies (Arthur *et al*. 2013). This device and related derived designs have been used in multiple studies (Shirangi *et al*. 2013, 2016; Philipsborn *et al*. 2014; Stern 2014; Coen *et al*. 2014; LaRue *et al*. 2015; Ding *et al*. 2016, 2019; Rezával *et al*. 2016; Clemens *et al*. 2018; O’Sullivan *et al*. 2018; Kerwin *et al*. 2020; Arthur *et al*. 2021; Shiozaki *et al*. 2022), although these devices incorporated several sub-optimal features.

To improve the design, reduce the cost, and allow even greater throughput, we completely redesigned a multi-channel device for recording courtship song and behavior. Our new design provides simultaneous audio and video recording of 96-channels. Video recording is critically important, because it can be difficult to resolve the social meaning of specific acoustic signals from audio signals alone. There has not previously been available a high throughput recording system that allowed simultaneous monitoring of audio and video. Our new system also incorporates a novel design for time-controlled, high-intensity LED illumination, allowing high-throughput optogenetic functional screens of labeled neurons.

## Results

The 32-channel recording device we designed previously (Arthur *et al*. 2013) had several limitations that we sought to address with a new design. Specifically, we sought to provide the following improvements.

1. Simplify assembly. The 32-channel system was complex and required extensive manual assembly steps.
2. Reduce maintenance. The 32-channel system used microphones attached with cables that are no longer manufactured and the cable connections fatigued and broke over time.
3. Lower apparatus cost: The National Instruments digitizer and MATLAB license required in the original design added substantial cost.
4. Provide an overall design that allowed end-users to flexibly re-organize the recording chambers to suit their requirements.
5. Simplify loading of flies into recording chambers while keeping males and females separated until experiment begins.
6. Improve the software user interface.
7. Provide synchronized camera frame triggers.
8. Record individual chamber temperatures.
9. Drive up to two optogenetic light sources.
10. Save all data and metadata to individual WAV files.
11. View summary of all 96 videos after completion of recording.
12. Compress individual videos.

We developed a recording apparatus that captures audio and video from 96 behavior chambers simultaneously. An 8 x 12 array of microphones records audio from each chamber from below and two cameras record video from above (Figure 1). All associated software is open source and available on Github (https://github.com/janelia-experimental-technology/FlySong). Flies are loaded into three 32 chamber plates (8 x 4 array) that sit on top of the microphones (Figure 2). The behavior chambers are built from mostly 3D printed parts and can be easily modified for novel designs. We have implemented designs that allow loading flies from the top, or from the bottom, and keep flies separated until the investigator is ready to initiate the recording session.

**Figure 1.**
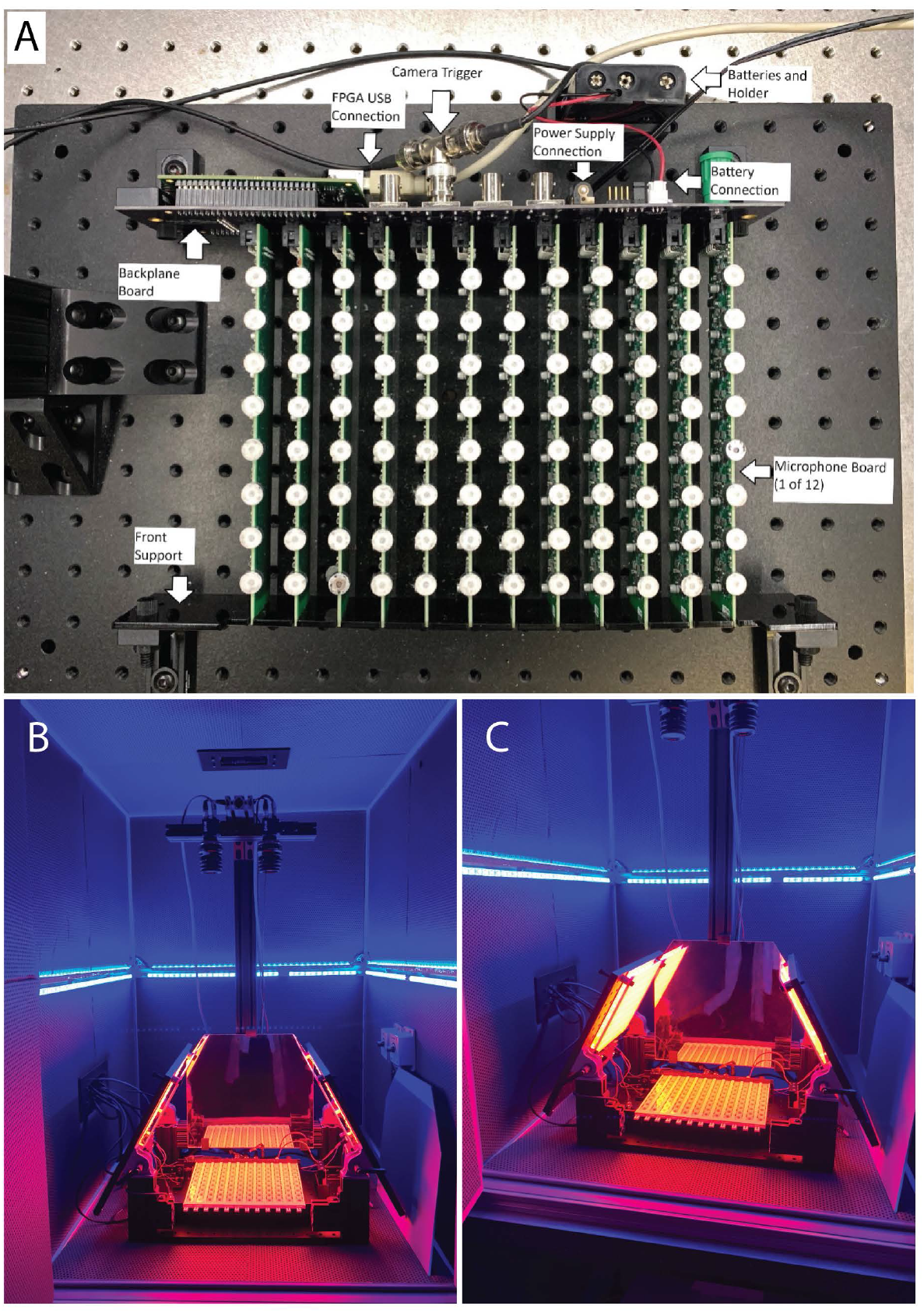
Hardware of the 96-channel *Song Torrent* audio recording rig. (A) Ninety-six microphones are arrayed on 12 microphone boards that are inserted into a single backplane board, which collects all microphone signals and sends the data stream to a computer via a USB connector and sends signals to trigger camera shutters. Behavior chambers are placed on top of microphones and are shown in Figure 2. (B) View illustrating the location and mounting of cameras. The rig is enclosed in a wind-proof box, which is itself mounted on an air table. (C) View illustrating illuminated LED panels with diffuser in front of lights.

**Figure 2.**
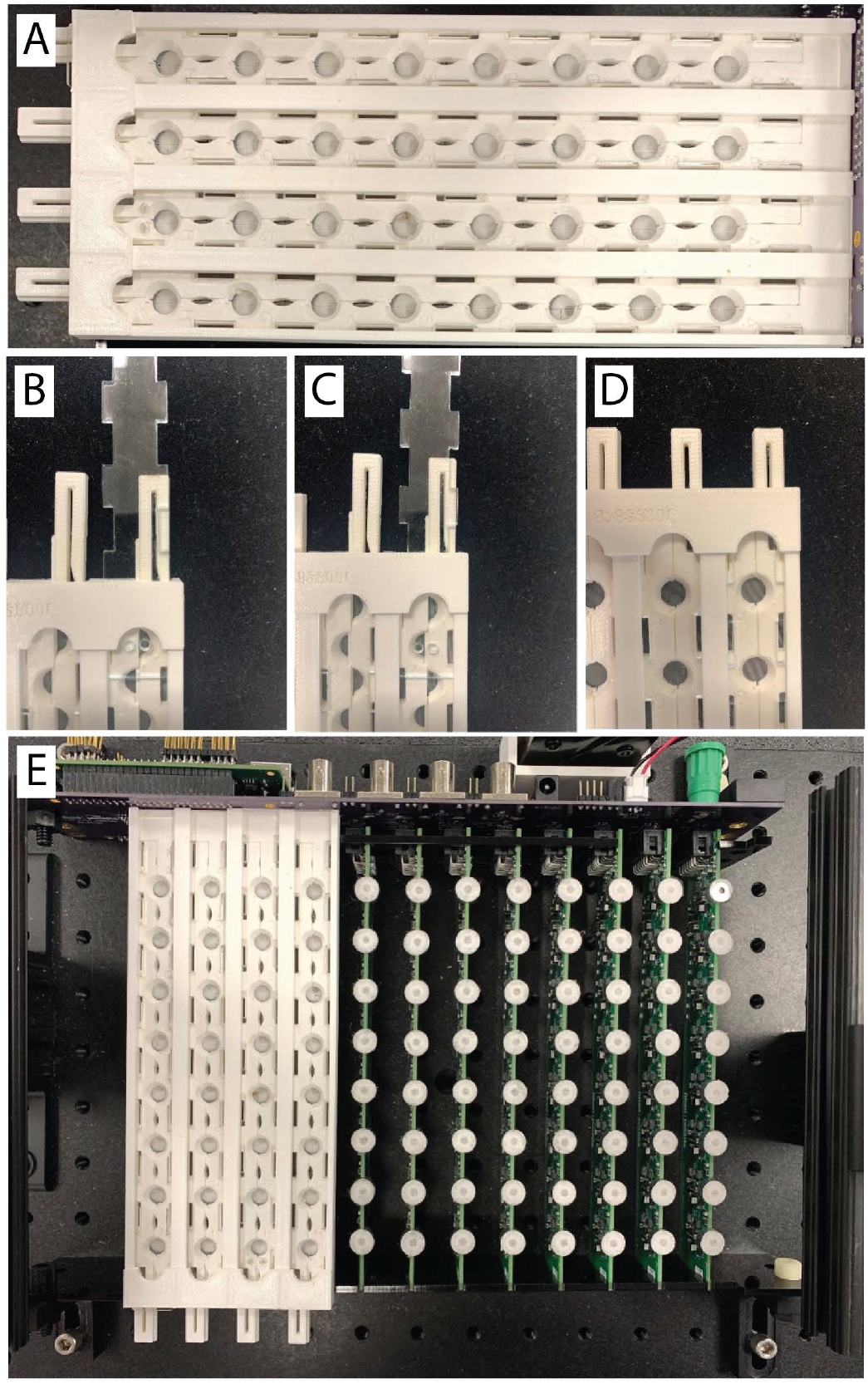
Behavior chambers for use with *Song Torrent*. (A) Top view of 32 channel behavior chamber panel. (B-D) Three views illustrating how the sliding mechanism of the behavior chamber can be used to load a pair of flies for a courtship assay. Initially the sliding mechanism is retracted so that each behavior chamber is a half circle (B, C). The top clear plastic slider is inserted so that one hole is positioned over one half circle (B). After the fly is loaded in the first half circle, the top plastic slider is moved so that a second fly can be loaded in the second half circle (C). This is continued until all required chambers are filled. After the 32-channel chamber is placed on top of microphones (D), the sliding mechanism is pushed in to form a full circle courtship arena (C).

The recording apparatus is composed of 12 printed circuit boards, containing 8 microphones each, that connect to a single backplane containing a Field Programmable Gate Array (FPGA) that provides an interface between the microphone boards and the computer (Figure 1A). If one board of 8 microphones fails, it can be replaced easily. The microphone boards can be modified for different layouts, depending on experimental requirements. Analog to digital conversion is performed on each microphone board, eliminating the need for an expensive separate analog-to-digital conversion unit.

The 3D-printed fly courtship chambers allow loading of flies initially into separate halves of each chamber, so that males and females remain separate until the experiment begins (Figure 2). The bottom of the chambers is a fine plastic mesh to allow sounds to be detected by microphones below. The top of the chambers is clear acrylic to allow video recording from above. We designed two different loading mechanisms: flies can be loaded through the top via holes in a specialized sliding cover or through the bottom, via slits cut into the white plastic mesh (Figure 2). In principle, the bottom loading design could allow automated robotic loading. To increase contrast of flies against the chamber background, so that video tracking software could be used to analyze fly behavior, we 3D printed chamber parts with white plastic, glued white mesh to the bottom of each chamber, and replaced the microphone covers with white mesh.

Video is captured by two high-resolution cameras. The provided software (*Fly Recorder*) separates the view of 48 behavior chambers captured by each camera into separate channels that are associated with each microphone channel. The software can be used to adjust the cameras position to capture higher resolution video from fewer chambers and to define the region of interest (ROI) for video capture. Using the provided software *Convert Video*, the individual videos can be compressed and converted from .mov to .avi format. Video files can be assembled into a single video using provided software (https://github.com/JaneliaSciComp/video_grid) for manual inspection of visual behavior across all chambers after completion of a recording session (Video 1).

Optogenetic excitation and inhibition of labelled neurons have become important assays for probing neural circuit function in *Drosophila*. We therefore designed high-intensity LED arrays that can be mounted onto the apparatus along with a mirror to redirect stray light (https://github.com/janelia-experimental-technology/RGB-IR-LED-Boards). This design provides consistent and strong illumination across a large area without obstructing the video recording during optogenetic manipulations. An example of video and audio recorded on *Song Torrent* during optogenetic stimulation of a fly expressing the red-shifted channel rhodopsin *CsChrimson* in the descending interneuron pIP10 that drives courtship song is shown in Video 2.

### Limitations of Song Torrent

We have identified several limitations with the current design of *Song Torrent*. First, while audio can be recorded without recording video, we have not currently implemented the ability to record video without recording audio.

Second, we have designed *Song Torrent* for study of *D. melanogaster* and closely related species of approximately the same size as *D. melanogaster*. Study of larger *Drosophila* species or of other insects may require redesign of the behavior chambers.

Third, we placed microphones sufficiently far apart to prevent crosstalk between channels when *D. melanogaster* and similarly sized *Drosophila* species sing courtship song. Study of louder acoustic signals, for example those produced by larger *Drosophila* species, may require redesign of the microphone spacing to prevent crosstalk between channels.

Fourth, optimal microphone recording requires that behavior chambers are placed precisely and directly on top of microphones. If the behavior chambers are displaced from the correct position, even by a few millimeters, the signal will be substantially reduced.

## Discussion

*Song Torrent* provides a high-throughput audio and video recording device that allows optogenetic stimulation and can be assembled from mostly off-the-shelf components. Most of the electrical components can be purchased using the design specifications that we have provided (https://github.com/janelia-experimental-technology/FlySong). Perhaps the trickiest step is to manually solder microphones onto the circuit boards. To facilitate construction, assembly, and use of *Song Torrent*, we provide a detailed assembly guide (Supplementary Material). We have found *Song Torrent* to be useful both for recording behavior of wild-type flies and for high-throughput screens of flies manipulated with optogenetic reagents (Shiozaki *et al*. 2022; Lillvis *et al*. 2023; Li *et al*. 2023).

We designed a specific layout of *Song Torrent* microphones and behavior chambers suitable for our experimental requirements, but all of these features can be altered prior to purchasing assembled components. In addition, video resolution can be increased by reducing the number of chambers recorded or by purchasing higher-resolution cameras. The flexibility of *Song Torrent* should allow audio recording of insects other than *Drosophila* species and of non-courtship related audio and visual behaviors.

## Supporting information

Song Torrent Technical Document and Assembly Instructions

Parts List for Song Torrent

Costs of Song Torrent parts

## Acknowledgements

We thank Mioara Andrea Gugiu for improving the layout of circuit boards, building boards, suggesting assembly techniques, and helping with the documentation. This article is subject to HHMI’s Open Access to Publications policy. HHMI lab heads have previously granted a nonexclusive CC BY 4.0 license to the public and a sublicensable license to HHMI in their research articles. Pursuant to those licenses, the author-accepted manuscript of this article can be made freely available under a CC BY 4.0 license immediately upon publication.

## Figure Legends

Video 1 – Example of output from the *video_grid* software, illustrating 96 pairs of courting flies from a single recording session.

Video 2 - Optogenetic activation on the Song Torrent apparatus of a fly expressing CsChrimson in the pIP10 descending interneurons drives courtship song in an isolated male. The male fly carries the stable split combination SS46542 (40556-p65ADZp [attp40]; 40347-pZpGdbd [attp2]) of Gal4 reagents driving 20XUAS-CsChrimson-mVenus [attp18] (Lillvis *et al*. 2023).

## Computer Code Availability

Software and custom hardware designs are available at Github via the following sites: https://github.com/janelia-experimental-technology/FlySong https://github.com/janelia-experimental-technology/RGB-IR-LED-Boards

## Supplementary Material

Song Torrent Technical Document.pdf: A PDF document providing technical specifications of the *Song Torrent* hardware and software and an assembly guide.

96ChannelRecordingRig.xlsx: An Excel file providing a full parts list for all components required to build a Song Torrent rig, excluding the computer, air table and wind-proof box.

Costs.xlsx: An Excel file providing the costs of *Song Torrent* components as of 2023.

## Notes

### Competing Interest Statement

The authors have declared no competing interest.

