## Supplementary material for "*Song Torrent*: A modular, open-source 96-chamber audio and video recording apparatus with optogenetic activation and inactivation capabilities for *Drosophila*": Song Torrent Technical Document and Assembly Instructions

### A 96 Chamber Fly Courtship Recording Array

### Technical Specifications

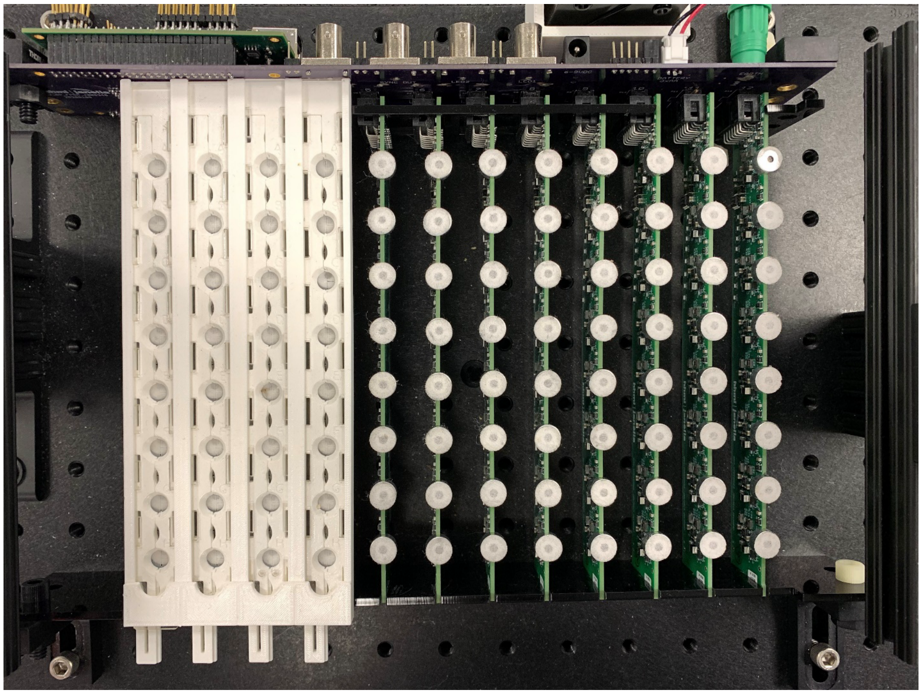

*Figure 1 - 96 Channel Recording Arena shown with one 32-channel behavior chamber assembly installed.*

#### System Overview

A 96 Chamber Array (Figure 1) and open-source software was designed to provide high throughput acquisition of *Drosophila* fly courtship songs. Up to 96 separate behavior chambers can be recorded simultaneously along with temperature data for each chamber. Behavior hambers are designed as three arrays of 32 chambers each and have sliding partitions to allow loading of separated males and females. The chambers can be top loaded using a mouth pipette through loading holes in a sliding cover or bottom loaded through a slit in the lower mesh. The latter method could be used by auto-loading systems. Two optogenetic light sources are provided and their timing and brightness can be controlled. A camera frame trigger is provided. The software allows any number and arrangement of chambers to be recorded. A separate time and brightness comma-separated variable (CSV) file defines the optogenetic light pattern. A separate CSV file can be used to predefine the individual sample names and their chamber locations, or this information can be entered interactively in the GUI and saved. All data (lights, temperatures, camera triggers, audio, sample information, etc.) are saved to .WAV files.

### Background on New Electronic Hardware Development

##### Audio Board

We designed a 96-channel array composed of modular components that allow easy replacement of faulty channels and alternate configurations. Wiring is minimized to reduce noise and complexity as well as to enhance reliability and ease of assembly. An array pattern of 8 x 12 was chosen with each eight-channel column built onto a single printed circuit board (PCB). The microphones are soldered directly to the PCB, but since they were not designed for automated assembly, they are added by hand after the rest of the board is assembled. Each microphone has its own signal conditioning circuit which is similar to the circuit used in Arthur et al. (2013) but upgraded to a trans-impedance design based on a Texas Instruments design (Caldwell 2015). It consists of microphone bias, amplification, and filtering. Surface mount components were used to minimize the size of the circuit. Audio circuit capacitors with film or electrolytic construction were used to provide optimal audio quality and eliminate possible microphonics (due to piezoelectric effect) that could be produced by ceramic capacitors.

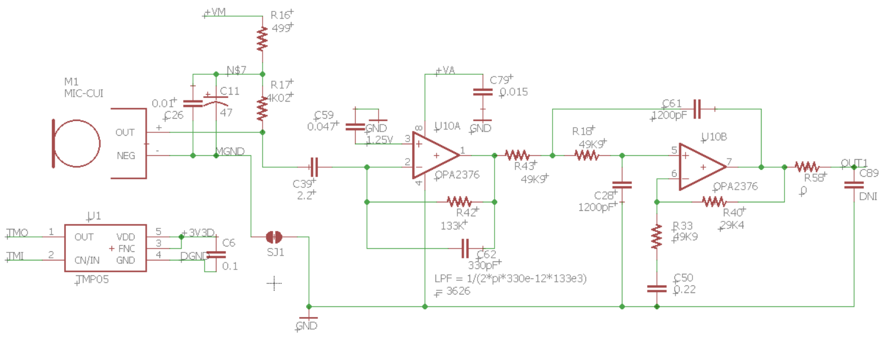

*Figure 2 - One channel of the microphone analog circuit, including temperature sensor*

The analog circuit is designed for single supply operation (Figure 2). Battery power (3 x 1.2V NiMH AA cells) is used to minimize power supply noise. This allows sufficient headroom to drive the 0-2.5 input of the Analog-to-Digital Converter (ADC). The first stage provides a gain of 634 (56dB) and the second stage provides an additional gain of 1.59 (4dB) plus a 2$^{nd}$ order low pass filter with a cutoff at 2658 Hz. Total gain is 1008 (60dB). The outputs from the eight amplifiers/filters are fed to an eight channel ADC with 16 bits of resolution.

The temperature sensors are digital devices that output a square wave with a duty cycle proportional to temperature. They are powered by an independent 3.3V low noise regulator to minimize adding noise to the microphone data. The period of each sensor is about 100 milliseconds. The sensors are daisy chained, so reading 96 sensors takes about 9.6 seconds. Note: all 12 boards must be attached to the backplane to record temperature data.

The signals are brought out to a 14-pin header to provide power as well as control and data I/O (all digital).

| Pin | Digital I/O Connector |
|---|---|
| 1 | ADC command in (MOSI) |
| 2 | ADC data out (MISO) |
| 3 | ADC clock in |
| 4 | ADC chip select |
| 5 | Temperature data in |
| 6 | Temperature data out |
| 7 | Temperature power enable |
| 8 | (no connection) |
| 9 | +5 volts in |
| 10 | +3.3V Analog Voltage |
| 11 | 5V Ground |
| 12 | Battery/Analog Ground |
| 13 | Battery/Analog Ground |
| 14 | Battery Voltage + |

**Backplane**

The backplane PCB holds one end of each of twelve microphone boards to support and space them as well as providing electrical connections. The microphone boards connect via the digital I/O connector. The backplane interconnects the twelve microphone boards to each other, a 3.6-volt battery supply, and the FPGA board. The board also holds the battery supply on/off switch and battery check circuit.

**FPGA**

The FPGA board provides the host computer connection via high-speed USB 2.0, three LED controllers, and the control and data signals to the microphone array via the backplane board. Due to the potentially high data rates (10,000 samples /second * 96 channels * 16 bits/channel = 15,360,000 bits/second), a Field Programmable Gate Array (FPGA) was used to provide the interface control functions between the host computer and the microphone array. A commercially available FPGA board (Opal Kelly XEM7100) was used as it simplified the FPGA board design and provided drivers and interfaces to simplify the host and FPGA software/firmware development. Three BNC connectors provide LED drivers that can control brightness via pulse width modulation (PWM). A fourth BNC provides the camera trigger output.

#### Assembly

The microphone boards should probably be built by an electronics assembly house due to the number of parts and their small size. A complete setup requires 12 microphone boards, with the cost per board at ~$300. The microphones are designed to be attached to wires but are instead soldered directly to the boards, so it will be necessary to hand solder the microphones after the circuit board is complete, as shown below:

Parts needed for soldering the microphones to the microphone board (Figure 3):

- Microphone heat sink
- Board support
- Board and eight microphones
- Solder wire
- Solder station and soldering pencil with gull wing tip
- Flux

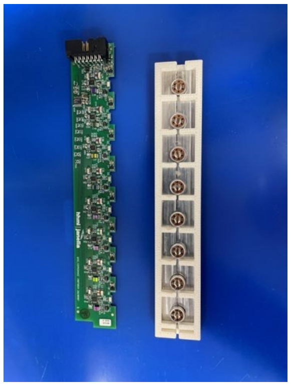

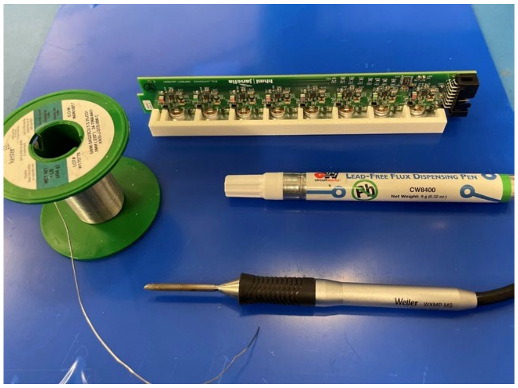

*Figure 3 – Parts used during assembly of a single 8-microphone board.*

The recommended solder tip for this procedure, if using Weller solder station, is T0054462499N (Figure 4).

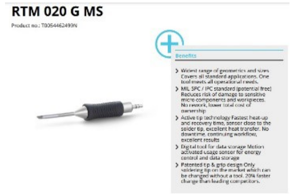

*Figure 4 – Recommended solder tip*

**Assembly Procedure**:

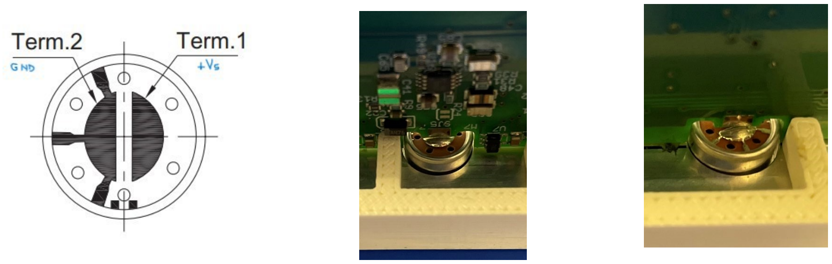

*Figure 5 – Diagram of back of microphone (left), and front (middle) and back (right) view of microphone soldered to channel.*

1. Note that microphones are polarized; terminal 2 is the ground side and has metal spokes that radiate out to the case (Figure 5). Place eight microphones in the heatsink so they are all aligned the same way with the terminals facing towards the long sides of the jig (Figure 3 & 5).

2. Place the board support on top followed by the assembled printed circuit board (Figure 6). The ground side of the microphones faces in the direction of the PCB that has no components. Align microphones centered under the solder pads on the PCB and in line with the board.

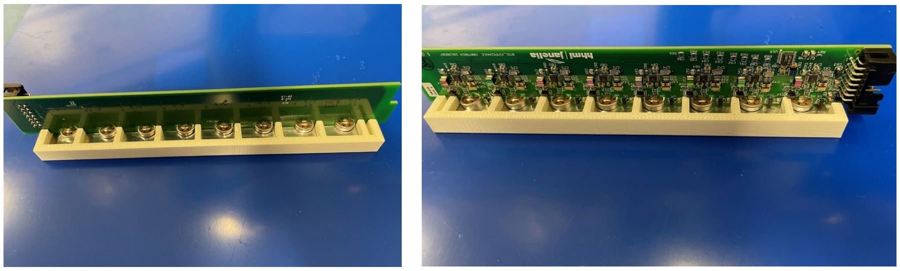

*Figure 6 – One 8-channel board placed in the board support for soldering of microphones.*

3. The microphones are then soldered (Figures 7 & 8):

   a. Wipe the connection point with a small amount of flux. Do not allow any fluid to enter the microphone.
   b. Add a substantial amount of melted solder wire to the solder pen tip.
   c. For a brief second, touch the microphone pad and board pad and remove it when the melted solder connects the two pads.
   d. Wipe clean with Isopropyl alcohol. Do not allow any fluid to enter the microphone.

4. If desired, the black covers can be replaced with white material. The covers are gently removed and, usually, the microphone has enough tack left on it to secure the new covers. The 'mic-covers.dxf' file can be used to laser cut covers.

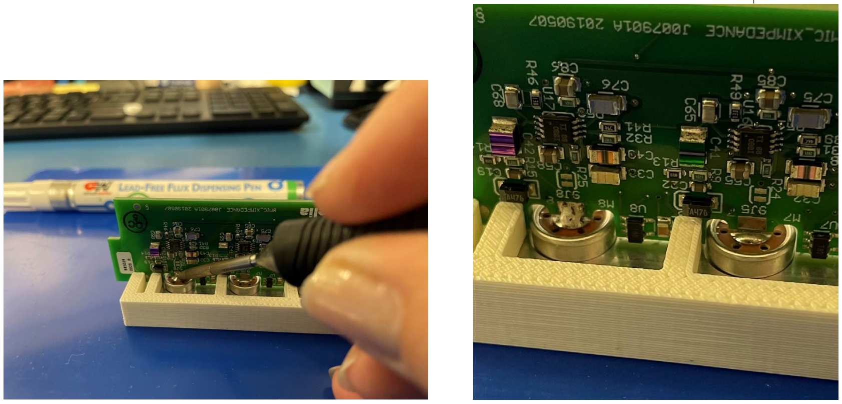

*Figure 7 – Photos of soldering left-most microphone (left) and of resulting soldered microphone to first channel (right).*

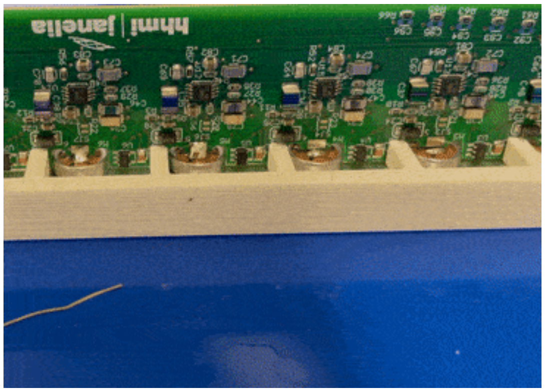

*Figure 8 – GIF illustrating how to solder microphone to channel.*

<u>Backplane Board</u>

The backplane board serves several functions: it mechanically positions the microphone boards, routes signals and power to the microphone boards, holds the FPGA board, buffers the USB data stream, and provides various outputs such as camera triggering and LED control.

Since only one board is needed per rig, it may be cost effective to hand assemble this board. This requires a bit of experience and the right tools such as a microscope and fine-top soldering iron (as shown above) or a reflow oven. The parts for this board (including PCB) cost approximately $300. If the assembled board is ordered from CircuitHub, the cost would be $850. The FPGA daughter board assembly is an additional $250.

Chambers

To provide an efficient mechanism for introducing male and female flies to each other as near to the start of recording as possible, we designed a new courtship arena array (Figure 9). Flies are first manually loaded into chamber halves. One half of each chamber is connected to a common slider that allows connecting the two halves just before the start of recording. The bottom of the array is covered by a mesh that can be cut with slits to allow bottom loading of the chambers (intended for automated loading). The tops of the chambers are covered with acrylic slides that have openings for manually loading flies from the top.

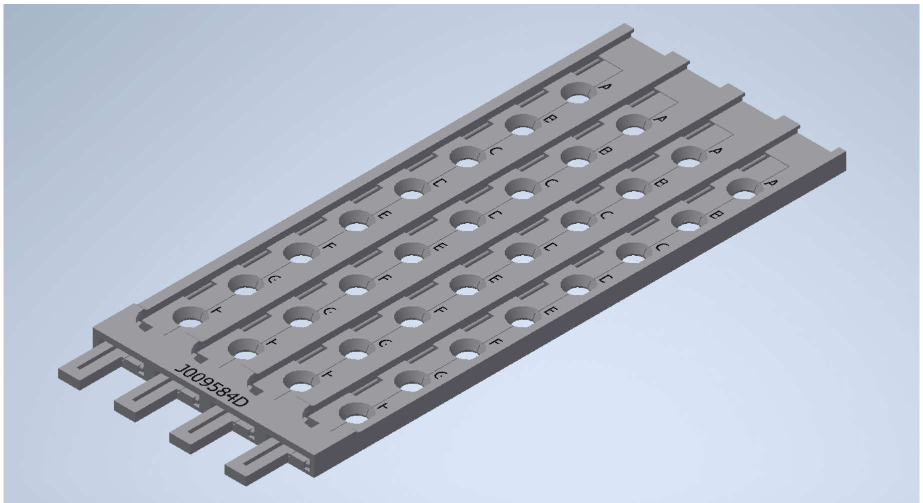

*Figure 9 – Diagram of 32-chamber arena base (without mesh or cover slides).*

The base of the array is 3-D printed out of ABS and forms the chamber base, the stationary chamber halves, the moveable chamber halves, and the open/close latch. The assembly contains 32 channels in a four by eight array so three of these assemblies are required to run all 96 channels. We print the chambers on a Stratasys F170 printer, using white ABS and 0.18 mm slice height.

The slides are laser cut from 1.6 mm clear acrylic. There are designs for 'loading' slides which have holes to pipette flies into the chambers as well as 'run' slides without holes. A loading slide can also be used to run an experiment.

The bottom of the chamber array is a mesh that is heat joined (e.g. with a soldering iron) to the bottom side of the chamber base (Figures 10 & 11). This mesh can be laser cut with T-shaped slits to allow either manual or robotic loading through the bottom of the chambers.  The mesh can be attached using a small soldering iron to melt the mesh to the base by quickly touching the soldering tip along the mesh at many points. Concentrate on attaching the mesh around each chamber. DO NOT weld the mesh to the sliding parts of the chambers! Also be sure to weld the mesh along the perimeters, especially along the one long thin edge. Have proper ventilation when performing this operation. The figures below are from an assembly using a black ABS base.

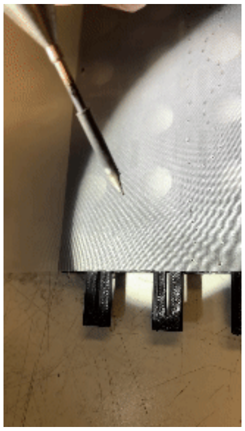

*Figure 10 – GIF illustrating how to attach mesh to back of courtship chambers.*

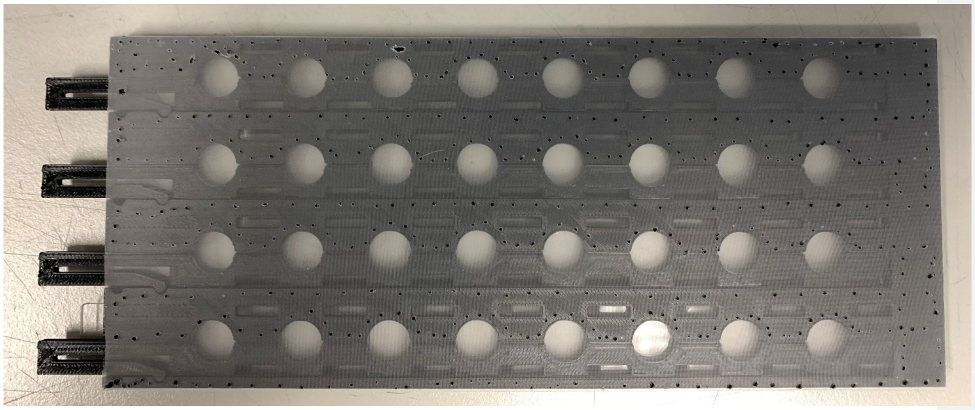

*Figure 11 – Mesh installed to back of courtship chambers.*

<u>Cameras</u>

Two FLIR Blackfly Monochrome cameras are used to provide detailed coverage of all 96 chambers.  A 50 mm lens with 50mm focal length is attached to each camera. The resolution of each camera is 5472 by 3648 pixels which provides about 1000 pixels per square mm at the behavior chamber.

Computer

A reasonably powerful computer with adequate RAM is required to handle the high data rates form the FPGA board and cameras. At a minimum, a USB 2.0 port is required for the FPGA board and two USB 3.1 connectors are required for the two cameras. Our current system has a computer with the following specifications:

- Processor: Intel Xeon W-2245 CPU @ 3.9 GHz
- RAM: 64 GB
- OS: Windows 10 Enterprise
- A large capacity solid-state drive is recommended to enable fast data transfers.

Rig Assembly

The components of the Fly Song Rig are assembled onto a 12" x 18" Thorlabs Breadboard. Supports for the backplane and microphone boards are cut from 0.062 acrylic. The rear support is fastened to the backplane and breadboard with right angle brackets. Two more brackets connect the front support to the breadboard. A battery holder is attached to the breadboard.

There are two assembly variations, depending on whether infrared/optogenetic illumination is needed:

For the basic setup, the cameras are mounted to a horizontal bar using 3-D printed brackets (Figure 12). The bar is connected to a vertical post using a right-angle bracket. The post is connected to the breadboard using two right angle brackets.

The setup that incorporates the LED panels requires adding two 7x7 RGB-IR panels, mounts, diffusers, and mirrors (Figure 13).

Mirrors are added to the back and front of the rig (front mirror removed to right side to show the rest of setup) to even out the lighting over the chambers. The uniformity of the light array is within 11% ((max-min)/max); measured with red color at 30% intensity (Figure 14).

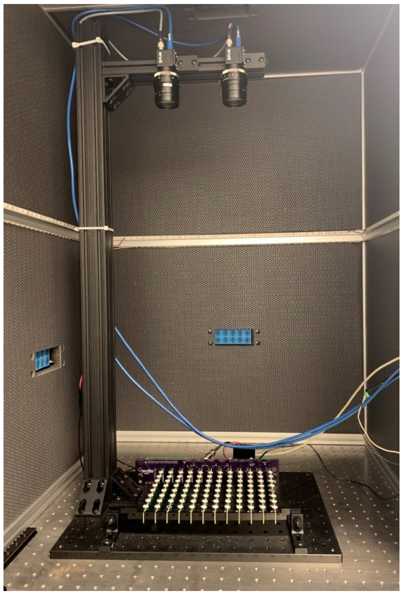

*Figure 12 – Fully installed audio recording rig placed on air table within wind-proof box.*

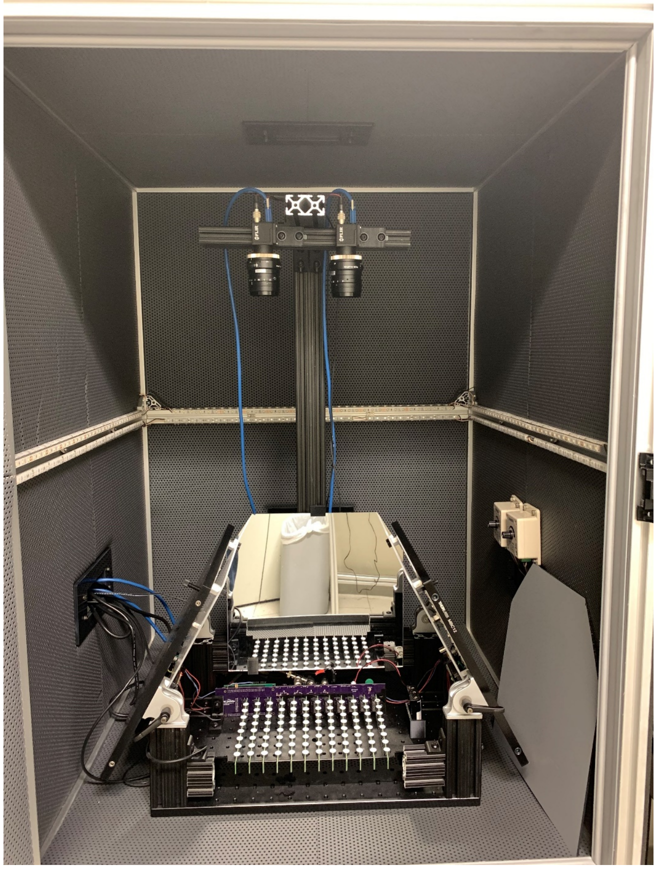

*Figure 13 – Fully installed audio and video recording rig with optogenetic light panels and mirrors. In this photo, the front mirror is removed and placed to the right.*

The two RGB-IR boards were designed at Janelia, each consisting of three sets of sixty-four LEDS placed uniformly over a 7” by 7” area. The three sets are generally assigned the primary colors red, green, and blue, but any available colors can be used. In addition, there are 256 infrared LEDs uniformly placed in the same area. Each color can be controlled independently. A diffuser helps to make the light pattern more uniform. The LED board is attached to a 12” by 12” Thorlabs breadboard for heatsinking and support. If high levels of light are needed for prolonged time, then one could substitute a liquid cooled breadboard to move heat outside the cabinet.  The breadboards are attached to locking pivots which

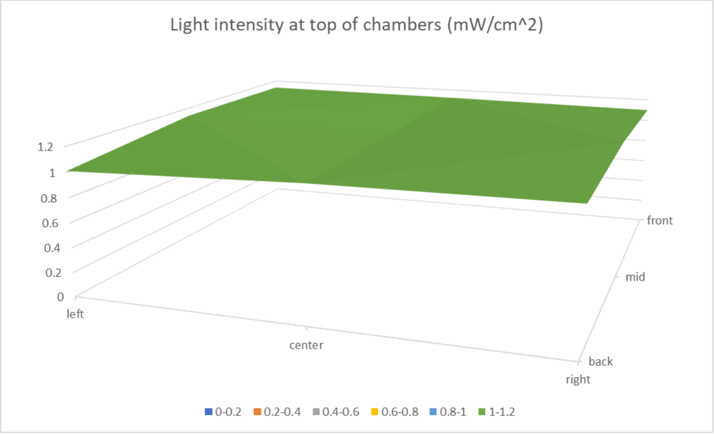

*Figure 14 – Measurements of optogenetics light intensity across the field of the entire 96-channel array.*

are, in turn attached to vertical posts. These posts are connected to the main breadboard using right-angle brackets. In this setup the camera post is attached to a cross beam attached to the rear verticals. A horizontal beam attaches to the top of the vertical post using a right-angle bracket and points toward the front of the rig. Underneath this bar is another bar mounted perpendicular. The cameras are mounted to this bar using 3-D printed brackets. The mirrors are cut from mirrored acrylic to fit between the LED boards in the back and front. The rear mirror can be permanently mounted to the structure, while the front mirror must be removable to allow loading and maintaining the rig. Small brackets cut from acrylic support the front mirror.

#### New System Hardware Setup

The Fly Song System consists of a backplane (Figure 15), up to twelve eight-channel microphone boards (Figure 16), a battery supply (Figure 17), wall AC-DC power supply (Figure 17), and behavior chambers (Figure 18):

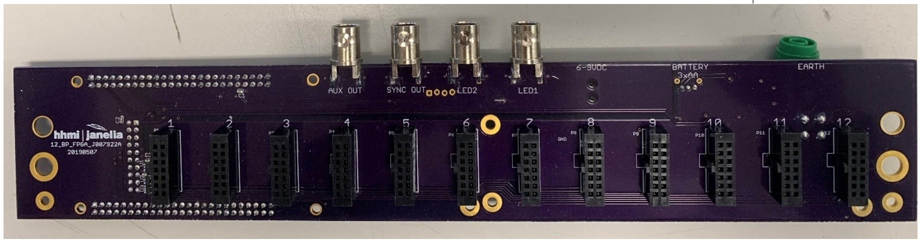

*Figure 15 – Front of backplane, showing the 12 slots for microphone boards.*

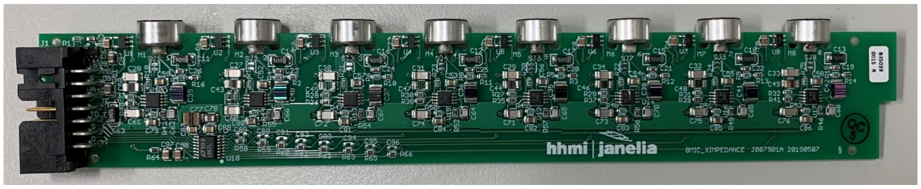

*Figure 16 – One fully assembled microphone board.*

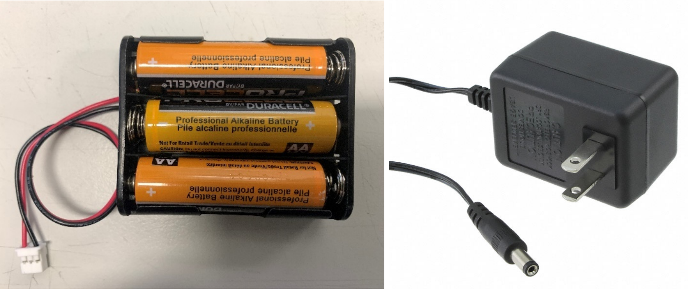

*Figure 17  - Battery back (left) and wall AC-DC power converter (right).*

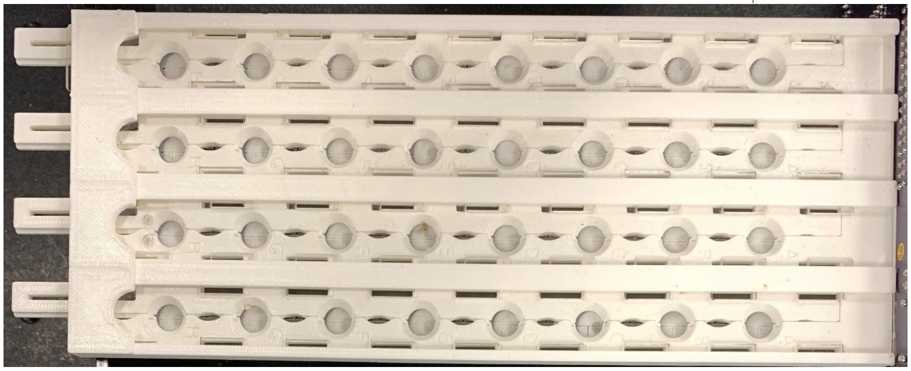

*Figure 18 – A fully assembled 32-channel courtship recording apparatus.*

Once the backplane and supports are mounted to the breadboard, plug the microphone boards into the backplane. Use one hand to support the back of the backplane while inserting each board (Figure 19). They can go in at a slight angle so that the front support can be left in place. The notch on each board should rest in the appropriate slot in the front support.

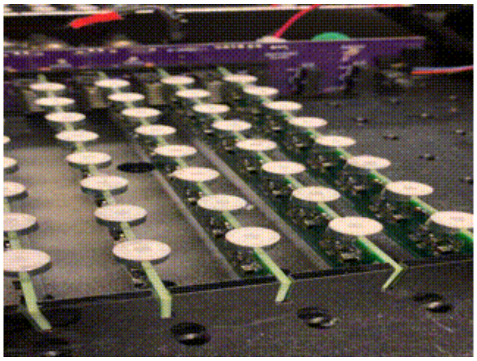

*Figure 19 – GIF illustrating how to insert a microphone board.*

Place three AA batteries into the battery holder, being mindful of the orientation (Figure 18). The battery holder can be placed in the plastic holder mounted to the breadboard. Plug the battery holder connector into the white connector on the back right of the backplane (Figure 20).

Plug the wall AC-DC power supply connector into the connector in the backplane. The connector faces down. Plug the power supply into a wall outlet.

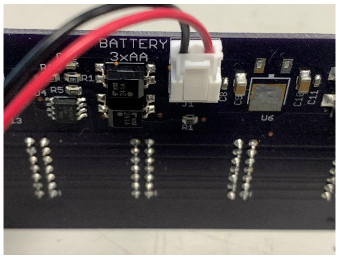

*Figure 20 – Photo illustrating connection of battery source to backplane.*

Plug a USB cable into the FPGA board (the smaller board plugged into the backplane). Plug the other end into your computer.

If camera triggering is required, connect the camera trigger cables to the Sync Out BNC on the backplane. For the cameras used in the original rig, trigger cables should be made up as shown in Figure 21.

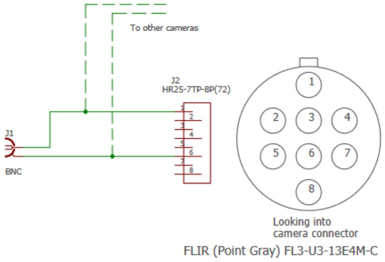

*Figure 21 – Circuit diagram of Sync Out BNC connector.*

A fully assembled rig is shown in Figure 22.

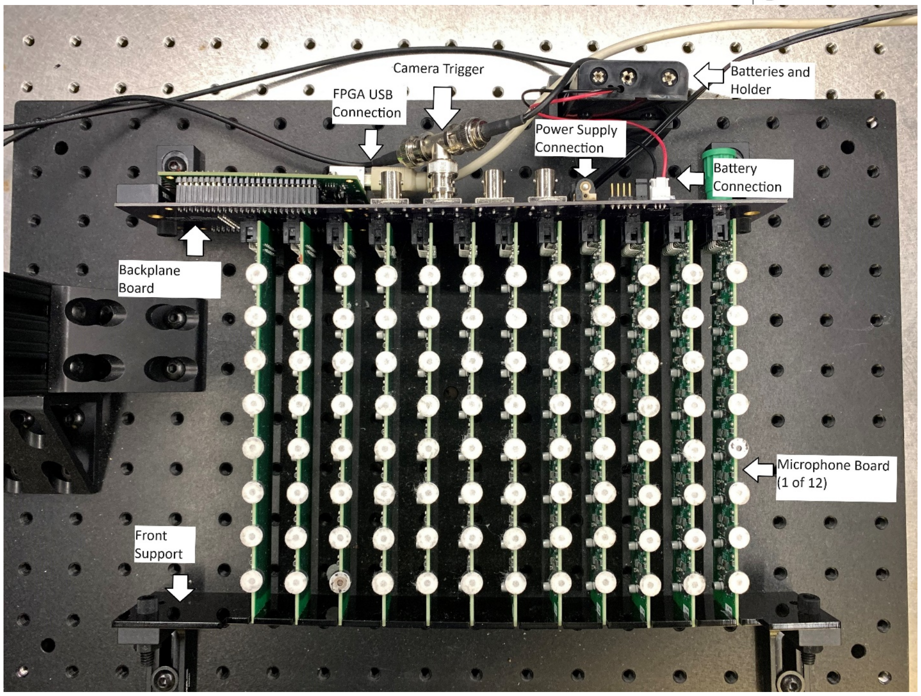

*Figure 22 – Top view of fully assembled 96-channel audio recording apparatus. Separate components are labelled.*

#### New System Software Setup

Camera Software

The camera software is available in the FlySongVideoInstall folder of the Github repository ([https://github.com/janelia-experimental-technology/FlySong](https://github.com/janelia-experimental-technology/FlySong)). Run "setup.exe" to install drivers and software.

FlySong Software

Do not plug the FPGA USB cable in until the software is installed.

Run 'FrontPanelUSB-DriverOnly-5.3.0.exe' to install the drivers for the FPGA board.

Unzip 'FlySong.zip' to the root directory ('C:'). You should end up with a new directory 'C:\\FlySong'. In that directory there will be example setup files and a Release directory. In that directory there will be an executable file named 'FlySong.exe'. This program provides the interface to the rig. A shortcut for this file can be placed on the desktop or task bar.

The FPGA board can now be plugged into the host computer.

When you double-click on the shortcut (or directly double-click on 'FlySong.exe') the software will load and recognize the FPGA board.

#### Setup and Running Experiments

**Fly Loading**

If using male-female pairs, slide the moveable parts of the chambers so that the chambers are separated into two half-moon sections (Figure 23). If using single flies with light activation, set the slides so that the chambers are whole circles. The slide has a slight indent to hold it in place. You may need to squeeze the lever on the tab at the end of the chamber slide to release the catch. For top loading, place a cover slip at the end of the chamber array away from the slide's tab and set the loading hole over the first half-chamber. Load the fly and move to the next half chamber. There are two holes that line up with each chamber half. If bottom loading, slide the cover slip all the way into place and load through the slits on the bottom of the chamber. If top-loading a single fly per chamber, then cover one of the loading holes to prevent escapes.

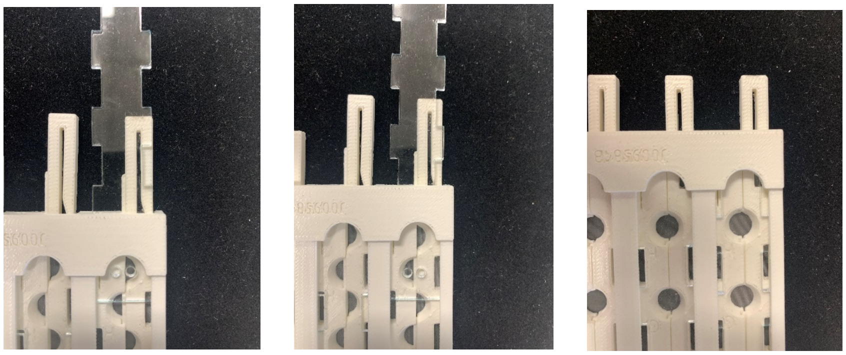

*Figure 23 – Three top views of courtship arena, illustrating how to use loading holes in sliding top to load flies sequentially into each half of each courtship chamber (left and middle) and with the top fully inserted to provide clear view of chamber and the two halves of each chamber slid into final position (right).*

Once flies are loaded, the chambers can be placed onto the recording rig (Figure 24). The chambers are set on top of the microphone boards with the tab end away from the backplane (Figure 24). Ascertain that the chambers are fully seated and centered over the microphones.

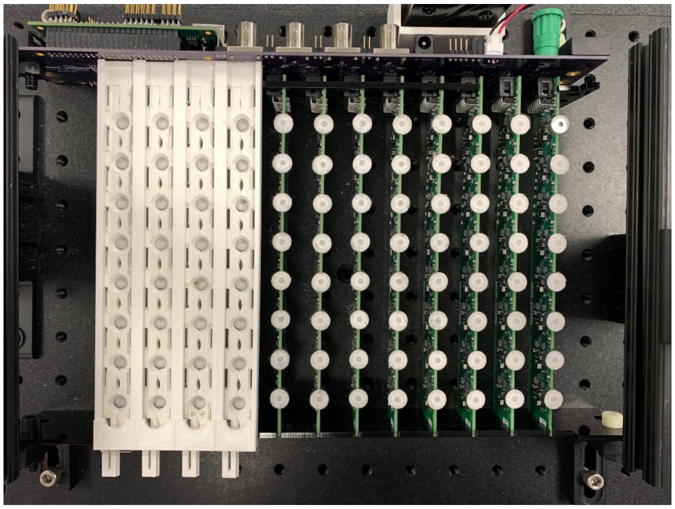

*Figure 24 – Top view illustrating the position of one 32-chamber courtship chamber loaded for recording.*

**Running an Experiment**

To run an experiment with optogenetic stimulation, a minimum of two files are required: an LED file and a Samples file. The LED file sets the timing and intensity of optogenetic lighting. The Samples file holds the identification and location of the flies in the rig.

<u>LED File</u>

The LED File is a CSV file with each line indicating the brightness and duration of up to three different light patterns. The first line must contain some text character beyond numbers or commas. The remaining lines control the LEDs. The first value is elapsed time in seconds. The next three values are brightness from 0 to 100 per cent (note, that although three LED brightness channels can be specified, only the first two are currently implemented in hardware). For example, to run an experiment with no lights for 5 seconds, then 30% brightness on channel 1 for 2 seconds, followed by no lights for 10 seconds, the file would look like this:

time, R, G, B
0, 0, 0, 0
5, 30, 0, 0
7, 0, 0, 0
17, 0, 0, 0

This experiment will run for 17 seconds.

If no light control is required, then the file consists of three lines: the first line with text followed by a line with 0 brightness, then a line with the total length of the experiment and zero brightness. To run an experiment with no LEDs for 10 second use:

time, R, G, B
0, 0, 0, 0
10, 0, 0, 0

These files are usually saved with the extension .TXT to make them easy to find. Though the software will automatically look for .TXT as well as .CSV files.

<u>Samples File</u>

The Samples File is a CSV file that holds the identifier for each chamber. The identifier can be any text, such as chamber number, genome, etc. This information can be entered before the experiment or in the Fly Song software. To create the Sample File offline, open a spreadsheet and enter the identifiers in the first twelve columns and eight rows of the spreadsheet. Unfortunately, the spreadsheet norm is to label columns with letters and rows with numbers, whereas the Fly Song rig uses letters for the rows and numbers for the columns, so disregard the spreadsheet numbering system. The upper left entry corresponds to chamber 1A, which is the rear chamber on the left-most microphone board (with the backplane at the rear – upper left in picture on first page). Note that the chamber letter designators are inscribed next to each chamber. Any chamber locations left blank will be assumed to be empty and data will not be stored for that chamber. The chambers can be loaded or empty in any manner desired; they do not have to be contiguous. An example with 4 flies loaded in various locations is shown below:

| | A | B | C | D | E |
|---|---|---|---|---|---|
| 1 | 1A Fly | | | | |
| 2 | | | | | |
| 3 | | | | | |
| 4 | | 2D Fly | | | |
| 5 | 1E Fly | | | | |
| 6 | | | | | |
| 7 | | | | | |
| 8 | | | | 4G Fly | |
| 9 | | | | | |

Save the file as a .CSV file. Do not add any other information in the spreadsheet.

One could also create the file using a text editor by entering the information using commas to delineate locations:

1A Fly,,,

,,,
,,,
,2D Fly,,
1E Fly,,,
,,,
,,,
,,,4G Fly

When loaded into the Fly Song program it would appear as in Figure 25. In the Fly Song application, the chamber information can be loaded and modified.

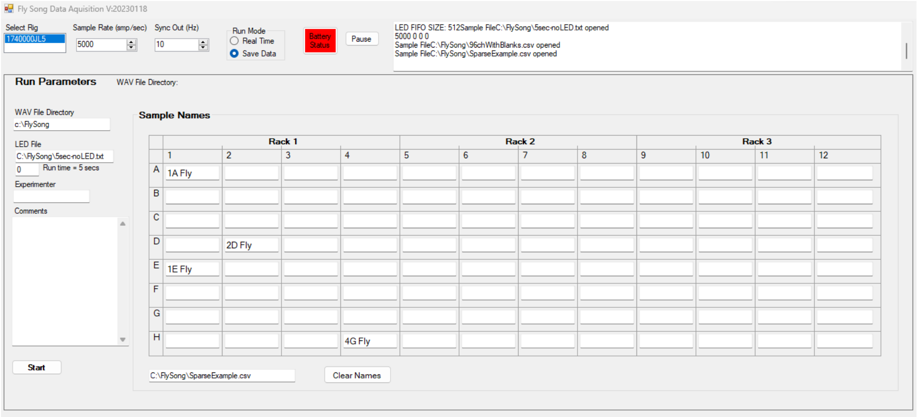

*Figure 25 – Screen shot of software illustrating the positions loaded for recording.*

**Fly Song**

Start up and Real Time

Start the Fly Song program. It will start up in real-time mode. This window will show the sampling setups and real time data from all 96 microphones. The screenshot in Figure 26 shows the microphones responding to whistling. A better test for the microphones is to use a signal generator app to generate a 200-300 Hz signal and run the phone speaker over each microphone. The upper left channel is 1A.

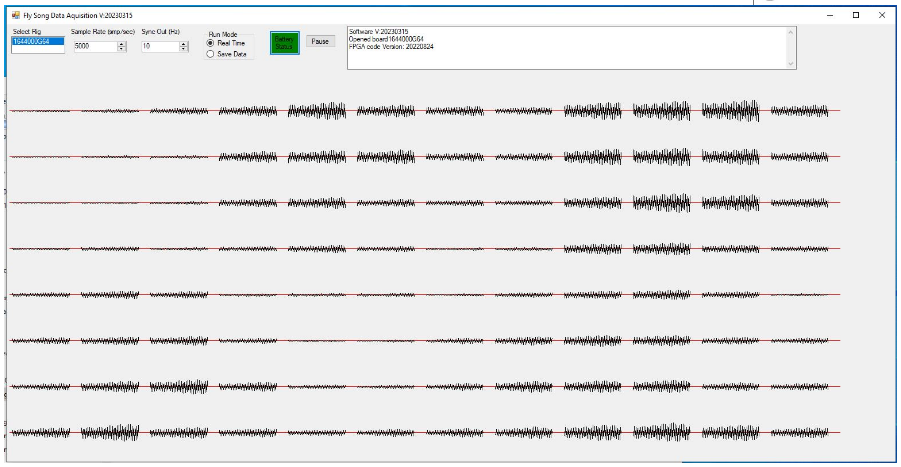

*Figure 26 – Screen shot of real-time mode showing signal detected by many microphones in response to whistling.*

The program will automatically find any attached rig(s) and display the serial number in the 'Select Rig' box. Generally, only one rig is attached, and, in that case, that rig is automatically selected and enabled. A status window at the right side of the screen displays status and debugging information.

The Sample Rate window displays the default sample rate. This can be modified to any value not exceeding 10,000 samples per second.

The Sync Out window displays the camera sync output rate. A square wave is sent to the SYNC BNC connector on the backplane at this rate. This can be used to trigger a camera to synchronize with the experiment. The output will be active in real-time mode. When switched to Save Data to run an experiment, the trigger will stop and restart when the experiment begins. This will synchronize the video to the start of the experiment.

Battery Status background will be red if the battery needs to be changed or charged. Click on it to check the status. It will turn green if the battery is ok.

The Pause button will pause the real time display.

Click on Save Data to open the window to run an experiment.

<u>Running an Experiment</u>

*Fly Song Data Acquisition (audio window)*

The Save Data window will show the top row of parameters, such as Sample Rate, etc. which can be modified (Figure 27). The real-time display is replaced by the Run Parameters.

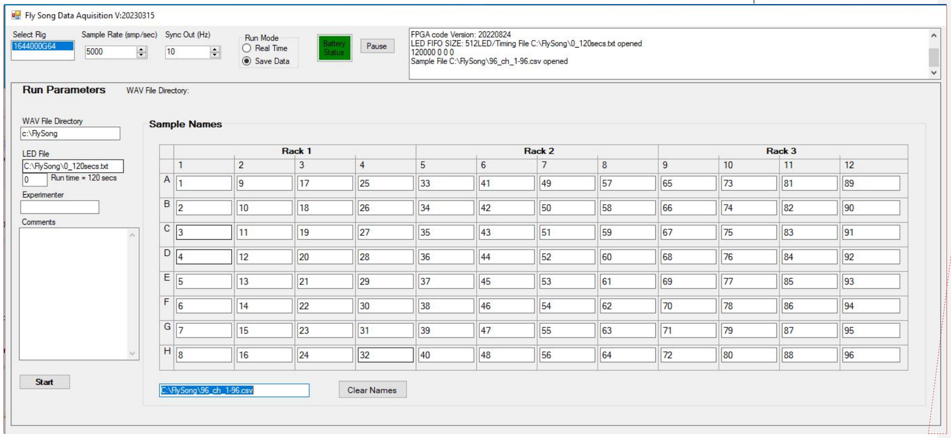

*Figure 27 – Screen shot of software illustrating audio recording parameters.*

**Commented [LJ1]:** I just noticed that there is a typo on how acquisition is spelled in the window (top left)

The default directory for storing data and setups is ‘C:\\FlySong’ but can be changed.

The WAV directory is where the data will be saved. All the files from a single run will be put into a subdirectory with the same base name as the files.

In the LED file field, browse to the desired LED file and load it. Under that field, the total run time will be displayed. When the experiment starts, the adjacent field will indicate elapsed seconds.

The Sample Names can be manually entered or loaded by clicking on the ‘Click to load Samples File’ field. This will bring up a browser to locate the file.

The experimenter can enter their name or other identifier into the Experimenter field if desired. This will be stored in the metadata for each WAV file.

If desired, comments can be added in the Comments field. This will also be saved into each WAV file.

Clicking on the Start button will start the experiment. Once started, a ‘STOP RUN’ button will appear to allow aborting the run. Otherwise, the experiment will run to the end and save the data to files.

Additional experiments can be run from this window, using the same parameters, or loading new ones.

*Fly Recorder (video window)*

Video recording is optional.

The run arrow button at the top of the window initiates the cameras in the window.

Parameters in the Exp Settings window (right) can be adjusted to optimize video quality.

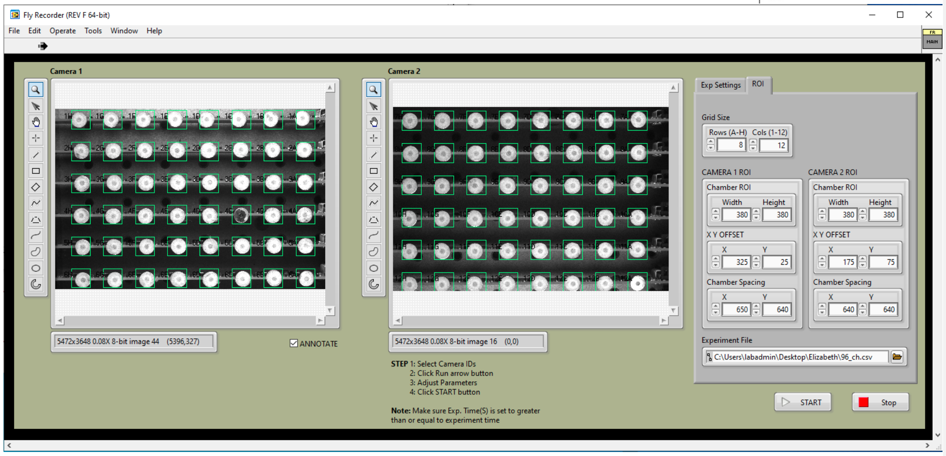

*Figure 28 Screen shot of software illustrating video recording parameters.*

Output Folder must be specified.

Parameters in the ROI window (right) can be adjusted to ensure that each chamber is captured by an ROI.

An experiment file (identical to the Sample File used in the audio window) must be indicated.

The START button readies the video recording. Subsequently starting the recording in the audio window triggers the video recording to start and synchronizes the audio and video recordings.

**IMPORTANT**: if recording video, this START button must be clicked first, followed by the Start button in the audio window with a relatively short delay to ensure that both recordings start and are synchronized.

#### Software and Firmware

All software and firmware is available on GitHub at:

**https://github.com/janelia-experimental-technology/FlySong**

##### FPGA Firmware

The FPGA firmware is written in Verilog using the Xilinx Vivado IDE and Opal Kelly libraries. Each microphone board has an eight-channel ADC with a Serial Peripheral Interface (SPI). The FPGA takes data from the array by selecting one channel on each of the twelve boards simultaneously. Only a single command line (MOSI) is needed since each board receives the same command (all get set to the same channel on their board). The FPGA reads the resulting conversion data on twelve separate input lines (MISO) for each board. The FPGA then converts the next channel. All channels are always converted, even if fewer than 96 are selected to be stored. The host can select which channels should be returned over USB. Simultaneously, the FPGA initiates temperature readings by sending a temperature reading start command pulse. When the duty cycle-based data is returned, the FPGA stores the pulse timing for each sensor. Actual temperature calculations are performed by the host software prior to saving the data. In addition, the FPGA acts on LED commands from the host and controls brightness by using pulse-width modulated (PWM) duty cycle to the LED drivers. The period of the PWM signal is 125 microseconds. In addition, a camera trigger output is available with the frame rate set from the host program. The three LED channels, camera frame trigger, and temperature channels are always stored. This allows the changes in LED brightness to be correlated to the data within one sample time. The LED brightness data is returned as a percent of full brightness. The temperature data is available at a much slower data rate and one channel at a time, but due to restrictions imposed by the WAV file format, all selected channels must be stored at the requested sample rate. For temperature, one channel is used to store which microphone position the data is being reported on (1-96), and the second channel is used to store the corresponding temperature in degrees Celsius * 100. At other times these channels are set to zero. The temperature data is also saved to a CSV file. Up to one hundred channels of data are transferred during an experiment. The first four channels are always LED brightness #1, LED brightness #2, temperature channel, and temperature data. The chosen audio channels are stored starting at channel 5. The FPGA logic, at the start of a new sample time, updates these four channels then goes into a loop where it converts all 96 analog channels. The first channel on each of the twelve boards is converted, followed by the second, etc. A 96-bit pattern holds a Boolean flag for each chamber denoting which data should be saved. The saved data is transferred to a 4096-word deep FIFO buffer. Other FPGA code coordinates with the host computer to transfer blocks of data to the host and receive commands from the host.

##### Host Software

The host software is written in C# using the Visual Studio IDE. This software provides a user interface to simplify setting up and running the microphone array and creating the WAV file outputs. To run the application, double click the Fly Song shortcut. The program should launch and display real time data for all channels. It will also show the selected rig (in case multiple rigs are attached), sample rate, camera trigger sync rate, and battery status (green indicates a good battery voltage). A status window indicates progress and any errors that may occur. The FPGA board must be connected via USB before the program is launched. The FPGA serial number should be shown in the status window. The rest of the window provides snapshots of all microphone channels in near-real-time, so the operator can verify all channels are working properly. Click on the 'Save Data' button to enter data recording mode. Here directories and file names for the various files required are selected (WAV file, LED control file, and, optionally, the samples file). The WAV file name is generated automatically and is in the following form: <year><month><date><"_"><hour><minute><second><"_"><channel>. The LED file is a CSV file with each line holding a brightness level (floating point 0-100) and the lights “on” time in seconds (floating point). The samples file is a CSV file consisting of microphone location (A1, etc.) and name of the sample on each line. This file is optional, because the sample locations and names can be entered into the spreadsheet built into the software (and can be then saved). Blank cells are assumed to be ‘off’ (not saved). The location is named by a letter denoting the channel on the board (lettered A-H, starting at the end closest to the backplane) and the number is the board number (1-12, starting at the left when the backplane is at the rear of the setup). The data is stored in memory during each run to minimize disk writing overhead and allow rearranging of data before saving to disc. The FPGA processes the ADC data from all twelve boards at the same time, cycling through the eight channels on each board and transmitting data to the host computer where it is saved in RAM. When the experiment concludes, each active channel is written out to a separate WAV file. Temperature, LED brightness, and camera trigger data are all saved to separate WAV files. Temperature is saved to a CSV file.

#### Data Files

Separate WAV files are created for each enabled microphone channel, each LED brightness channel, and the camera frame trigger data. The temperature data, being asynchronous to the audio channels, is saved in two WAV files: one for temperature channels and one for the temperature data corresponding to each channel.  Since the temperature data has many redundant entries, a CSV file is created that removes the redundant data and stores the time (in milliseconds form the experiment start) of a temperature reading, the channel, and the temperature in degrees Celsius. In addition, the information displayed in the status window of the FlySong program is stored to a log file.

**Data Format**

The microphone data is in signed 16-bit little-endian format.

The camera frame trigger is represented as an increasing modulo-2^6 value.

LED brightness is in percent brightness x 10.

Temperature in the Wave file is stored as degrees C x 10.

Temperature in the CSV file is stored as degrees Celsius.

**File Naming**

All files produced in single run share the same file name beginning:

YYYYMMDD_hhmmss_

Where:

- YYYY = year

- MM = numeric month
- DD = date
- hh = 24-hour clock hour
- mm = minute
- ss = second

(Underlines separate parts of the name)

Wave files and video files add a channel descriptor and the '.WAV' or '.mov' extension, respectively.

**Commented [LJ2]:** Added a bit about the video files. Not sure if anything else useful is required. I think it would be useful to include the Convert Video code and a short description.

- name = channel name in the form of:
  - row/column_ordinal channel number, i.e. the first channel is 1A_1 and the last is 12H_96
  - LED1 – LED 1 brightness values
  - LED2 – LED 2 brightness values
  - SYNC – frame trigger count
  - TEMPCH – temperature channel
  - TEMPVAL – temperature values

The log file adds 'log.txt' to the base name.

The temperature CSV file adds 'TEMP.CSV' to the base name.

#### WAV File Format

The Wave files are designed to store the data as well as the metadata to fully document the experimental conditions. The standard WAV file format structure is used so that third-party software, such as MATLAB and Audacity can display as much of the metadata as possible. Different WAV file reader programs have varying abilities to decode the metadata structures. Some metadata was required to be entered into less used (but still standard) information structures provided by the WAV format. Standard WAV file format information can be found on the internet.

The file is organized as follows:

**RIFF Chunk Descriptor**

- RIFFnnnnWAVE: standard opening WAV file header, 'nnnn' is the number of bytes in the file following this header.

**Fmt Sub-Chunk**

- Sub chunk 1 size
- 1: for audio file format (PCM)

- Number of channels: number of enabled microphones + 4 for the LED and temperature data
- Sample rate
- Byte rate
- Block Align
- Bits per sample (16)

**data Sub-Chunk**

- Sub chunk size
- Data – the data is consecutive 16-bit values in little-endian format.
- Note: the non-microphone channels are stored as separate WAV files with data:
  1. LED brightness over time. Saved as percent brightness percentage * 10 (0-1000)
  2. Camera Sync – starts at 0 and increments every trigger, modulo 2^6
  3. Temperature channel (1-96)
  4. Temperature value (degrees C * 100)

**INFO LIST Sub-Chunk**

- nnnn : size of subchunk from end of 'INFOLIST'
- INAM : this holds the file names of the LED file (required) and Samples file (optional) "LED=LedFileName SMP=SamplesFileName" or "LED=LedFileName SMP= (no file)"
  Where LedFileName and SamplesFileName are the actual names of the respective files
- IPRD : Serial Number of FPGA board
- ISFT: Software versions for the host and FPGA code, both in YYYYMMDD format:
  "FlySongDAQ:YYYYMMMDD FPGA:YYYYMMDD"
- IART: Name of experimenter as entered in the host software
- ICMT: Comments added by the experimenter in the comment box
- ISBJ: Channel name:
  Fly song channels: location and name entered in the channel spreadsheet
  LED channel: channel (LED1 or LED2) and "%BRIGHT"
  Camera trigger file: "SYNC,FPS"
  Temperature channel file: "TEMP,CH"
  Temperature value file: "TEMP,DEG C"

#### Temperature CSV File Format

Temperature data is encoded into two WAV files, for channel number and value. It is also saved as a CSV file that may be more convenient. This file has three columns: milliseconds from start of experiment, channel, and temperature in Celsius. The first row of the file has the column names:

| milliseconds | channel | temperature (degC) |
|---|---|---|
| 99 | 1A | 31.15 |
| 203 | 1B | 30.35 |
| 310 | 1C | 29.99 |
| 415 | 1D | 29.91 |
| 513 | 1E | 30.2 |
| 607 | 1F | 30.63 |
| 712 | 1G | 29.82 |

The remaining rows hold the actual data. The temperature sensors are daisy chained, so the data is entered in serially. Therefore, each reading is taken at a slightly different time. It takes about 10 seconds to read all 96 channels. All channels are always read, regardless of the Sample Names file.

<u>Recommended components</u>:

Wind-proof box— The microphones used in Song Torrent are highly sensitive to the particle velocity component of audio and therefore detect air displacement. Song Torrent does not require a sound-proof box. Instead, a wind-proof box can provide sufficient insulation against air displacement that would be detected by the microphones.

Air table — We have found that building vibrations and resonant sounds can be transmitted to microphones because the microphones are, effectively, shaken through the air. We therefore recommend placing the Song Torrent apparatus on top of an air table that can efficiently dampen vibrations above approximately 50HZ. Cables connect Song Torrent to a computer, which is not kept on the air table. It is possible that cheaper solutions, such as placing Song Torrent on a partially inflated bicycle tire, may be sufficient to dampen building vibrations to retain the characteristic SNR of Song Torrent.

Parts List and Files:

See ‘96ChannelRecordingRig.xlsx’ at https://github.com/janelia-experimental-technology/FlySong for a tabbed spreadsheet of lists of all parts, code, and manufacturing files.

Printed Circuit Boards were designed in Eagle PCB and are available at https://github.com/janelia-experimental-technology/FlySong/tree/main/Electrical.

Mechanical parts were designed in Autodesk Inventor and plans are available at https://github.com/janelia-experimental-technology/FlySong/tree/main/Mechanical.

PC Software was developed in C# under Visual Studio and is available at https://github.com/janelia-experimental-technology/FlySong/tree/main/FlySongSource.

FPGA Firmware was developed in Verilog using the Xilinx Vivado IDE and Opal Kelly libraries and is available at https://github.com/janelia-experimental-technology/FlySong/tree/main/FPGAsource/FPGAwSRAM.
